# Nutritional quality and genetic differences of five amaranth cultivars revealed by metabolome profiling and whole-genome sequencing

**DOI:** 10.1101/2025.03.03.641344

**Authors:** Khalid Sedeek, Andrea Zuccolo, Umair Ali Toor, Krishnaveni Sanikommu, Sunitha Kantharajappa, Luis Riveraserna, Rod A. Wing, Magdy M. Mahfouz

**Author notes:** Correspondence: Magdy Mahfouz.

## Abstract

**Background:** Amaranth (*Amaranthus* spp.) has high nutritional quality, with edible grain and leaves, and many agronomic advantages, making it a promising part of the solution for global food insecurity. However, we lack comprehensive metabolomic and genome sequence data for many cultivars. To support the improvement of this versatile, sustainable crop, a detailed metabolome profiling of the edible grains and leaves and genome sequencing resources is required for the widely cultivated grain amaranth cultivars such as Coral Fountain (CF), Emerald Tassels (ET), Golden Giant (GG), Hopi Red Dye (HR), and New Mexico (NM).

**Results:** Through a non-targeted high-throughput metabolic profiling using ultra-performance liquid chromatography-tandem mass spectrometry, we precisely determined the whole-grain and leaf metabolites of these five cultivars. This analysis identified 426 and 420 metabolites with known chemical structures in the grain and leaf, respectively. The five amaranth cultivars differed significantly in the levels of several nutritionally valuable compounds in grains and leaves, including sulfur amino acids, vitamins, and chlorogenic acids, as well as potentially anti-nutritive compounds, such as oxalate and raffinose family oligosaccharides. On average, the cultivars CF and ET had more favorable levels of most identified health-promoting compounds compared to GG, HR, and NM. In addition, we provide high-quality reference genome sequences for the five cultivars using the PacBio Sequel II sequencing platform with an estimated genome size of 465–483 Mb comprising 46.9–48.7% repetitive elements. We generated an iso-seq library from different amaranth plant parts and utilized it to predict the amaranth genes and annotate their function into their respective gene ontology terms.

**Conclusions:** These resources will assist in breeding improved amaranth varieties and identifying targeted genes for trait modification and advancement through genome editing and engineering technologies.

## Background

Emerging needs to enhance food security and agricultural sustainability are driving researchers to look at alternative crops that produce high-nutrition foods and are adapted to adverse growth conditions. For example, amaranth (*Amaranthus* spp., Amaranthaceae family) is a genus of flowering plants comprising approximately 85 species and 400 varieties that are widely distributed in temperate and tropical regions throughout the world [1, 2]. Unlike many grain and vegetable crops, amaranth produces edible seeds and leaves. The seeds are commonly eaten as a cereal grain, ground into flour, popped like popcorn, cooked into porridge, or germinated into nutritious sprouts [3, 4]. Amaranth grain is a good source of protein and is rich in several essential amino acids [5, 6]. The protein content of the grain varies between 14% and 18.4% depending on the variety, surpassing that of wheat (*Triticum aestivum*), maize (*Zea mays*), and rice (*Oryza sativa*). In addition, amaranth grain contains notable quantities of fat (5–9%), starch (57–62%), and dietary minerals, such as iron, calcium, magnesium, manganese, phosphorus, and copper [2, 6–9]. Furthermore, it contains beneficial lipids, a high fiber content, and substantial levels of health-promoting bioactive compounds such as antioxidants. Its lack of gluten makes amaranth grain a good alternative for people with celiac disease and/or gluten sensitivity [8, 10]. In addition, amaranth leaves can be cooked similarly to spinach (*Spinacia oleracea*).

Amaranth has many favorable agronomic characteristics, such as rapid growth, the ability to thrive in poor soil conditions, and remarkable adaptability to adverse climatic conditions and plant diseases. Moreover, its C_4_ photosynthesis allows it to efficiently use light and water [7, 11]. Cultivated amaranths are currently mainly used for grain and as a vegetable; however, other amaranths are used as forage, ornamental plants, and in the cosmetics and dye industries [12–14]. The most economically important species, domesticated mainly for grain production, are prince’s feather (*Amaranthus hypochondriacus*), red amaranth (*A. cruentus*), and pendant amaranth (*A. caudatus*). They produce small grains with diverse colors and nutritional attributes. Their leaves vary in size and color (green or purple) and can be used for food and feeding livestock. These nutritional and agronomic attributes collectively position amaranth as a promising alternative crop for the future.

Producing improved amaranth cultivars will require researchers to develop genomic resources and produce a comprehensive analysis of the nutrient composition across diverse amaranth cultivars to identify nutrient-rich genotypes suitable for targeted trait engineering and agricultural development. Metabolomics approaches have been widely applied to uncover the chemical composition of several cereal crops [15–18]. Here, we conducted a non-targeted high-throughput metabolic profiling of five popular amaranth cultivars using ultra-performance liquid chromatography–tandem mass spectrometry (UPLC–MS/MS) to accurately identify and quantify whole-grain and leaf metabolites. Through in-depth comparisons among cultivars, we aimed to identify the cultivar with the richest grain and leaf contents of health-promoting chemicals. Additionally, although the chloroplast genome has been sequenced in many amaranth species [13, 19, 20], the nuclear genome of most cultivated varieties remains largely unexplored. Thus, we sequenced the whole genome of the five cultivars to provide valuable sequence resources essential for gene discovery, genetic variation analysis, and the improvement of desirable traits through genome editing and breeding technologies.

## Results

### Untargeted metabolomic profiling identifies significant differences among grain and leaf metabolites in the five cultivars

To investigate the nutritional quality of amaranth grain (_G_) and leaf (_L_) tissue, we determined and compared the metabolic profiles of five cultivars belonging to the three main grain-amaranth species: *Amaranthus caudatus* ‘Coral Fountain’ (CF) and ‘Emerald Tassels’ (ET); *A. cruentus* ‘Golden Giant’ (GG) and ‘Hopi Red Dye’ (HR); and *A. hypochondriacus* ‘New Mexico’ (NM) (See representative photographs of panicles, grains, and leaves in Fig. 1A-C). We identified 496 compounds in the grain (426 known and 70 unknown; Table S1). We identified more compounds (559) in the leaf tissues than in grain (496); however, 139 of these 559 compounds did not match known identities in the library (Table S2). We determined that the compounds of known identity are involved in nine super-metabolic pathways and 59 sub-pathways. These nine super-metabolic pathways comprise amino acids, carbohydrates, lipids, CPGECs (cofactors, prosthetic groups, and electron carriers), nucleotides, peptides, phytohormones, xenobiotics, and secondary metabolites. The abundance of all identified compounds gives an indication of the nutritional value and the physiological state of the grain and leaf. To assess the differences in metabolite composition among amaranth cultivars, we performed Welch’s two-sample *t*-test; the number of compounds in each pairwise comparison that met the statistical cut-off (*p* ≥ 0.05) as well as the directionality of the change are summarized in Table 1. For instance, the comparison of CF_G_ to NM_G_ resulted in 230 significantly differentially abundant compounds, with 170 and 60 compounds with a higher abundance in CF_G_ and NM_G_, respectively. We observed broad differences between samples belonging to different groups; we visualized the compound classes that relate to these differences as heatmaps using scaled imputed data (Fig. 2A, B; Table S3 and S4). The pattern seen in the heatmap supported the high reproducibility among the triplicates for all samples. Other observations of significance included the higher median response (inferred by the prevalence of red color in most of the compound classes) for many compounds in the grains and leaves of cultivars CF and ET compared to the other comparisons (Fig. 2A, B). This effect occurred in most of the general classifications (amino acids, lipids, cofactors, and secondary metabolites). Moreover, we visualized the differences in each metabolite abundance in the grains and leaves of the five cultivars as a boxplot in Table S5 and S6.

**Fig. 1.**
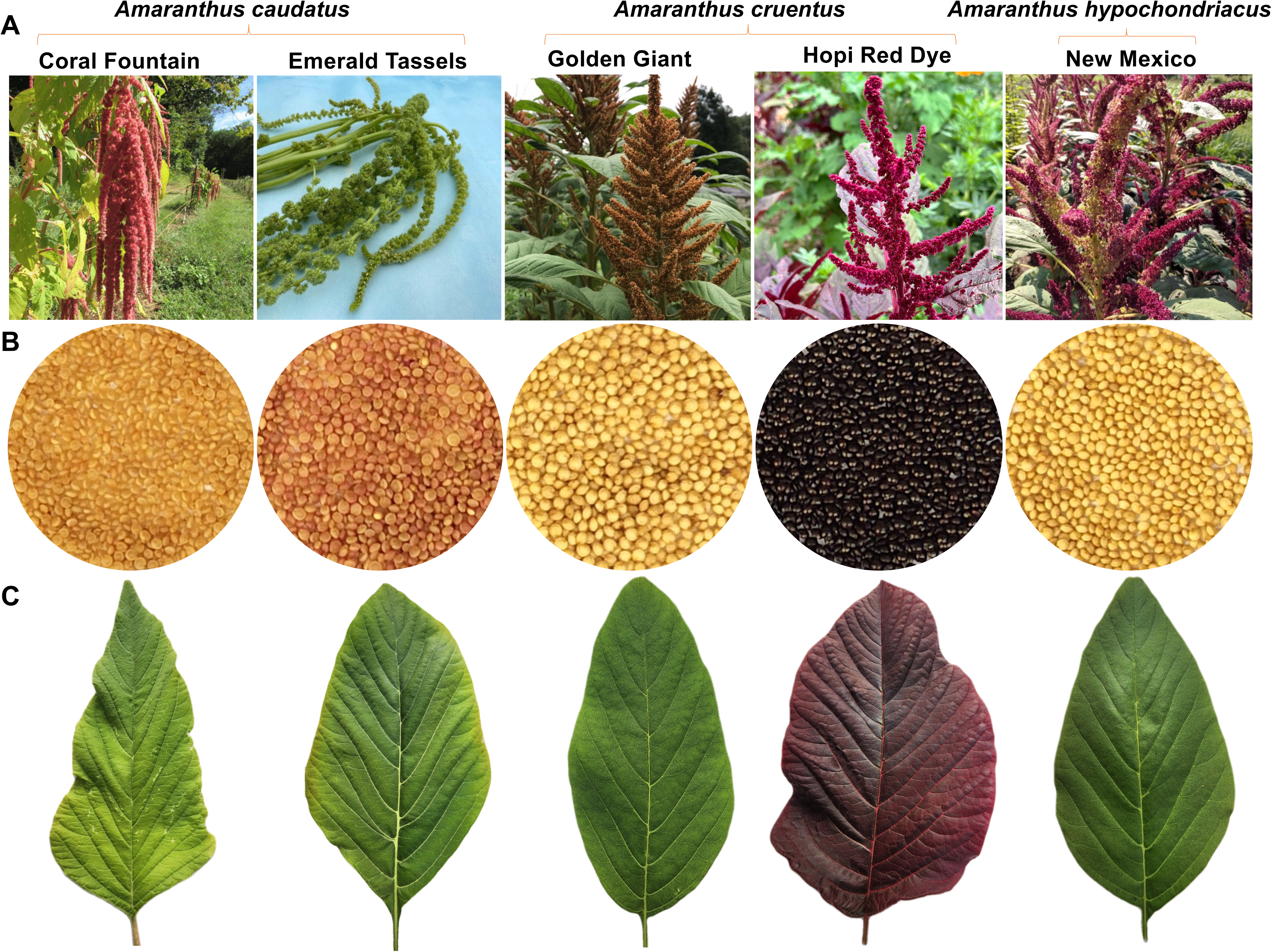
Representative images of the five cultivars. A) panicle (image provided by Hudson Valley Seed Company, United States), B) grain, and C) leaf.

**Fig. 2.**
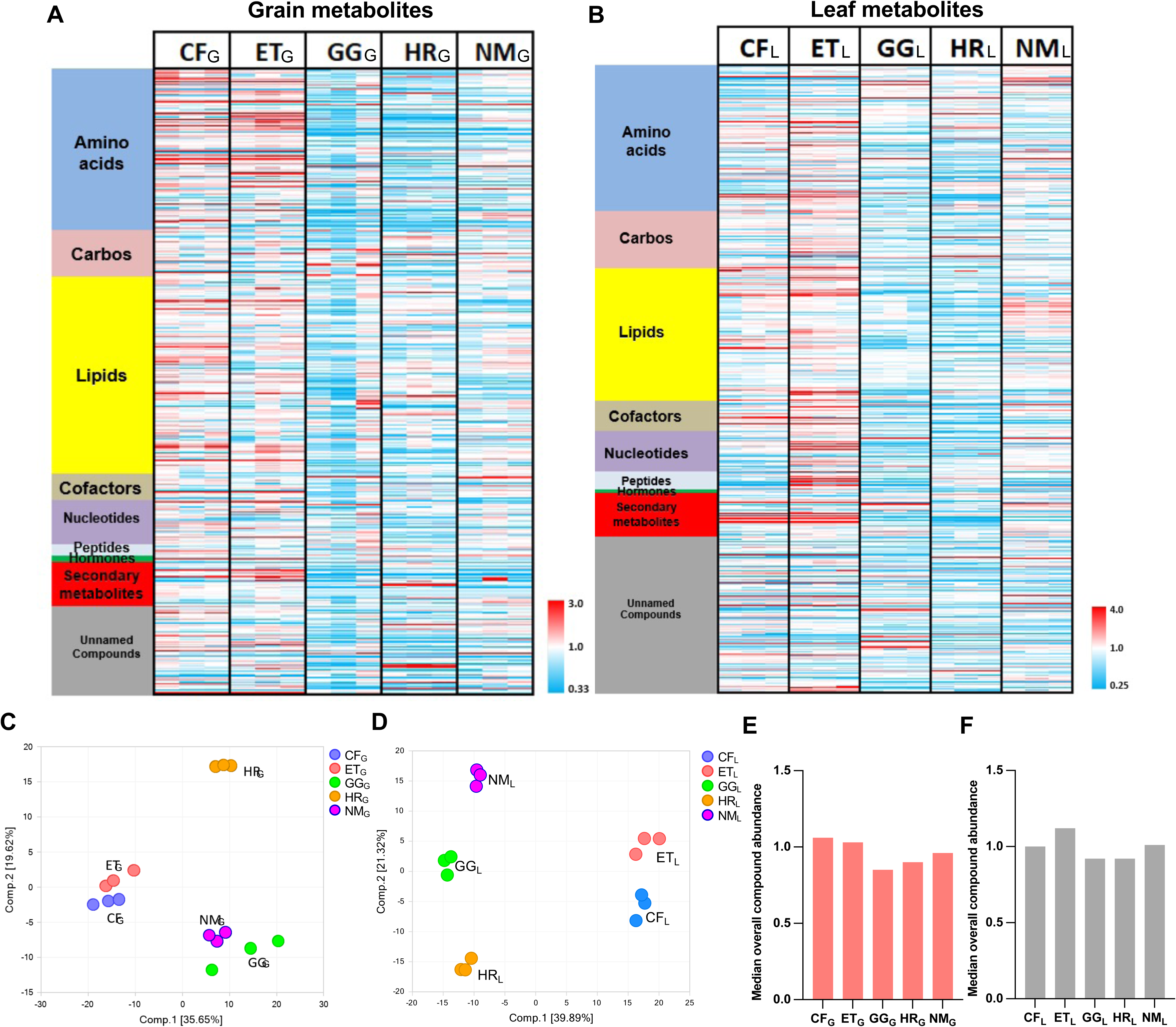
Metabolome screening of the grain and leaf of five amaranth cultivars and their overall content variation. A) Heatmap visualization of the contents of grain metabolites. B) Heatmap visualization of the contents of leaf metabolites. Each column represents one sample, with three replicates were analyzed per cultivar. Red indicates levels higher than the median value across the column, while blue indicates levels below the median. White indicates values near the median. C) PCA plot for grain metabolites. D) PCA plot for leaf metabolites. Positive scores along principal component 1 or 2 indicate higher levels of the identified compounds, while negative scores indicate lower levels of the identified compounds. E) Median overall compound abundance in grains of the different cultivars. F) Median overall compound abundance in leaves of the different cultivars. CF, Coral Fountain; ET, Emerald Tassels; GG, Golden Giant; HR, Hopi Red Dye; NM, New Mexico. (_G_), grain; (_L_), leaf.

**Table 1.** Summary of compounds meeting the statistical significance cut-off (*p* ≤ 0.05) in pairwise comparisons of amaranth grains and amaranth leaves of the five cultivars and relative direction of change. The number of more abundant compounds is shown in blue, and the number of less abundant compounds in red.

| Tissue | Cultivar ratio | Total compounds<br>$p \leq 0.05$ | Compounds<br>↑ \| ↓ |
| --- | --- | --- | --- |
| Grain | CF<sub>G</sub> / NM<sub>G</sub> | 230 | 170 \| 60 |
|  | ET<sub>G</sub> / NM<sub>G</sub> | 214 | 147 \| 67 |
|  | GG<sub>G</sub> / NM<sub>G</sub> | 60 | 16 \| 44 |
|  | HR<sub>G</sub> / NM<sub>G</sub> | 208 | 78 \| 130 |
|  | ET<sub>G</sub> / CF<sub>G</sub> | 128 | 62 \| 66 |
|  | GG<sub>G</sub> / CF<sub>G</sub> | 194 | 37 \| 157 |
|  | HR<sub>G</sub> / CF<sub>G</sub> | 258 | 60 \| 198 |
|  | GG<sub>G</sub> / ET<sub>G</sub> | 183 | 41 \| 142 |
|  | HR<sub>G</sub> / ET<sub>G</sub> | 244 | 60 \| 184 |
|  | HR<sub>G</sub> / GG<sub>G</sub> | 127 | 68 \| 59 |
| Leaf | CF<sub>L</sub> / NM<sub>L</sub> | 361 | 191 \| 170 |
|  | ET<sub>L</sub> / NM<sub>L</sub> | 340 | 210 \| 130 |
|  | GG<sub>L</sub> / NM<sub>L</sub> | 274 | 84 \| 190 |
|  | HR<sub>L</sub> / NM<sub>L</sub> | 316 | 115 \| 201 |
|  | ET<sub>L</sub> / CF<sub>L</sub> | 266 | 170 \| 96 |
|  | GG<sub>L</sub> / CF<sub>L</sub> | 353 | 126 \| 227 |
|  | HR<sub>L</sub> / CF<sub>L</sub> | 344 | 120 \| 224 |
|  | GG<sub>L</sub> / ET<sub>L</sub> | 368 | 97 \| 271 |
|  | HR<sub>L</sub> / ET<sub>L</sub> | 358 | 87 \| 271 |
|  | HR<sub>L</sub> / GG<sub>L</sub> | 297 | 146 \| 151 |

### Principal component analysis separated the cultivars based on their grain and leaf metabolic profiles

To assess similarities and differences between the metabolic profiles of the grain and leaf from the five cultivars, we performed principal component analysis (PCA), one-way analysis of variance (ANOVA) tests, and Tukey’s multiple comparisons. Cultivars were clearly separated in the PCA plot based on their metabolite contents (Fig. 2C, D). The first component (x-axis) differentiated CF and ET from the three other cultivars (Fig. 2C, D), which was driven by many compounds related directly or indirectly to vitamins or secondary metabolites (Fig. S1 and S2). In the case of the grain metabolome, the content of galactinol, the precursor to raffinose family oligosaccharides (RFOs), was low in CF_G_ and ET_G_ but high in the three other cultivars (Fig. S1). *Alpha*-tocopherol showed a similar pattern, while *delta*-tocopherol and tryptophan followed an opposite trend (Fig. S1). The second component (y-axis) differentiated GG, HR, and NM as well as separated ET from CF (Fig. 2C, D). This separation was driven by compounds that differed among groups within the two classes created by component 1. We noticed that GG_G_ and NM_G_ are similar in their compound abundance pattern, while HR_G_ appeared strongly separated, as exemplified by the contents of the phenylpropanoid compounds ferulate and 4-hydroxycinnamate (*p*-coumarate) (Fig. S1). We observed the opposite pattern (with HR_G_ showing low abundance) in the contents of the auxin catabolite indole-3- carboxylate and stachydrine, suggesting the involvement of high-level regulatory pathways differentiating the cultivars (Fig. S1). In the case of the leaf metabolome, quinate and feruloylquinate contents were high in CF_L_ and ET_L_ and very low in GG_L_, HR_L_, and NM_L_ (Fig. S2). Conversely, shikimate and kaempferol-glycoside contents were very low in CF_L_ and ET_L_ but higher in GG_L_, HR_L_, and NM_L_ (Fig. S2). The contents of the membrane phospholipid 18:2/18:2 glycerophosphaethanolamine (GPE) and the carotenoid beta-cryptoxanthin differentiated these groups, favoring ET_L_ and NM_L_ (positive scores and high on the y-axis (Fig. 2D); the negatively scoring stachyose and tryptophan fit a near mirror image of these plots, favoring CF_L_ and HR_L_ (Fig. S2).

### Cultivar-related pathway variation

When we calculated the average abundance of all named compounds within the five major compound classes in each amaranth cultivar (Fig. S3), we observed that the relative patterns largely follow the same pattern as the median overall response, as seen in Fig. 2E-F. Indeed, CF and ET tended to have generally higher metabolite accumulation tendency relative to the other cultivars, but the magnitudes of differences between the cultivars were more pronounced, especially in the contents of amino acids, lipids, cofactors, and secondary metabolites (Fig. S3). The very high average secondary metabolite abundance in CF_L_ and ET_L_ (up to 300-fold or more for some compounds (Fig. 6 I-J) compared to the other cultivars appeared to be highly significant in terms of cultivar-specific pathway expression, and potentially high nutritional properties. Specifically, with respect to nutrition, amaranth grain proteins are high in methionine and lysine. We also detected several other sulfur-containing compounds, such as methionine-related metabolites (S-adenosylmethionine [SAM], S-adenosyl-L-homocysteine [SAH]), as well as cysteine and its metabolites, in amaranth grain and leaf tissues (Fig. S4, Table S1).

Methionine levels in amaranth grain cultivars varied by less than a two-fold difference among cultivars, with CF_G_ and ET_G_ having slightly higher amounts (Fig. S4B). Notably, the highest relative levels of free methionine in leaves occurred in CF_L_, with NM_L_ being the lowest (Fig. S4J). The content of the catabolite 5-methylthioribose was especially high in CF_L_ and ET_L_ (Fig. S4C). This compound is an intermediate in the methionine salvage pathway (Fig. S4A), which recovers organic sulfur. Another important function of the methionine salvage pathway is to maintain low levels of 5-methylthioadenosine (MTA), which is an inhibitor of polyamine biosynthesis, a critical pathway for cell division and growth. MTA is produced from SAM, a metabolically important and central compound that provides methyl groups for a wide array of biochemical reactions across all major pathways, as well as donating carbon backbones for the production of ethylene, polyamines, and the important siderophore nicotianamine.

We also observed differences among cultivars for the contents of several compounds in the leaf metabolome associated with plant stress mitigation (Fig. S5A-F). These compounds included glutathione, which is a key metabolite related to oxidative stress, and several quaternary ammonium compounds (e.g., glycine betaine, stachydrine, trigonelline), which are important for relieving osmotic stress [21]. We noticed a high association among the quaternary ammonium compounds, all being highest in CF_L_. The siderophore nicotianamine is important for the transport of divalent cations in the phloem, and is known to confer tolerance to metal toxicity [22]. Cultivar differences among these various stress-related compounds, while mostly not directly related to nutrition *per se*, may have agronomic importance, as one of the key traits making amaranth a desirable crop is its ability to resist drought and other adverse environmental conditions [23].

### Vitamins are abundant in amaranth grain and leaf tissues

One class of metabolites with direct dietary value are vitamins essential for human nutrition. Fig. 3 displays the contents of several vitamins, or their related metabolites, present in the amaranth grain and leaf tissues. We observed variation in the contents among the grains of the five amaranth cultivars for all these compounds, which included several B-vitamins (B1, B5, and B6), nicotinate (a precursor for NAD^+^ biosynthesis), two forms of E vitamins (*alpha*- and *delta*-tocopherols), 4-oxo-retinoate (a compound with high vitamin A activity), and gulonate (a precursor for vitamin C) (Fig. 3A–H). We observed no clear pattern that would indicate one cultivar as being generally nutritionally superior to the others. For example, HR_G_ was highest in pyridoxine (B6) content but tended to be lower in the content of most other compounds, while NM_G_ was the richest in pantothenate (B5), *alpha*-tocopherol, and 4-oxo-retinoate but very low in several others. Interestingly, the contents of *alpha*-tocopherol and *delta*-tocopherol appeared to mirror each other in the cultivars, with the *alpha* species being lowest in CF_G_ and ET_G_ but high in GG_G_, HR_G_, and NM_G_, while *delta*-tocopherol showed the opposite pattern (Fig. 3E–F). The pathways responsible for the biosynthesis of these compounds diverge at the point of the direct precursor for *delta*-tocopherol, which forms the *delta* species via catalysis by a cyclase enzyme. To produce *alpha*-tocopherol, two methylation steps are required, one before the cyclization step and one after; we thus speculate that the differences observed may be related to the methylation potential in different cultivars. Amaranth leaves showed detectable levels of vitamin C, NAD^+^, riboflavin, and additional metabolites of pyridoxine (Fig. 3I–P). Because we detected many relevant compounds in leaves, we ranked the cultivars with respect to their tendency to produce higher levels of the various vitamin metabolites. We scored compounds based on the order of their abundance (1 = highest, 5 = lowest), then averaged across the compounds (Table S7). ET_L_ had the highest average ranking, followed by CF_L_.

**Fig. 3.**
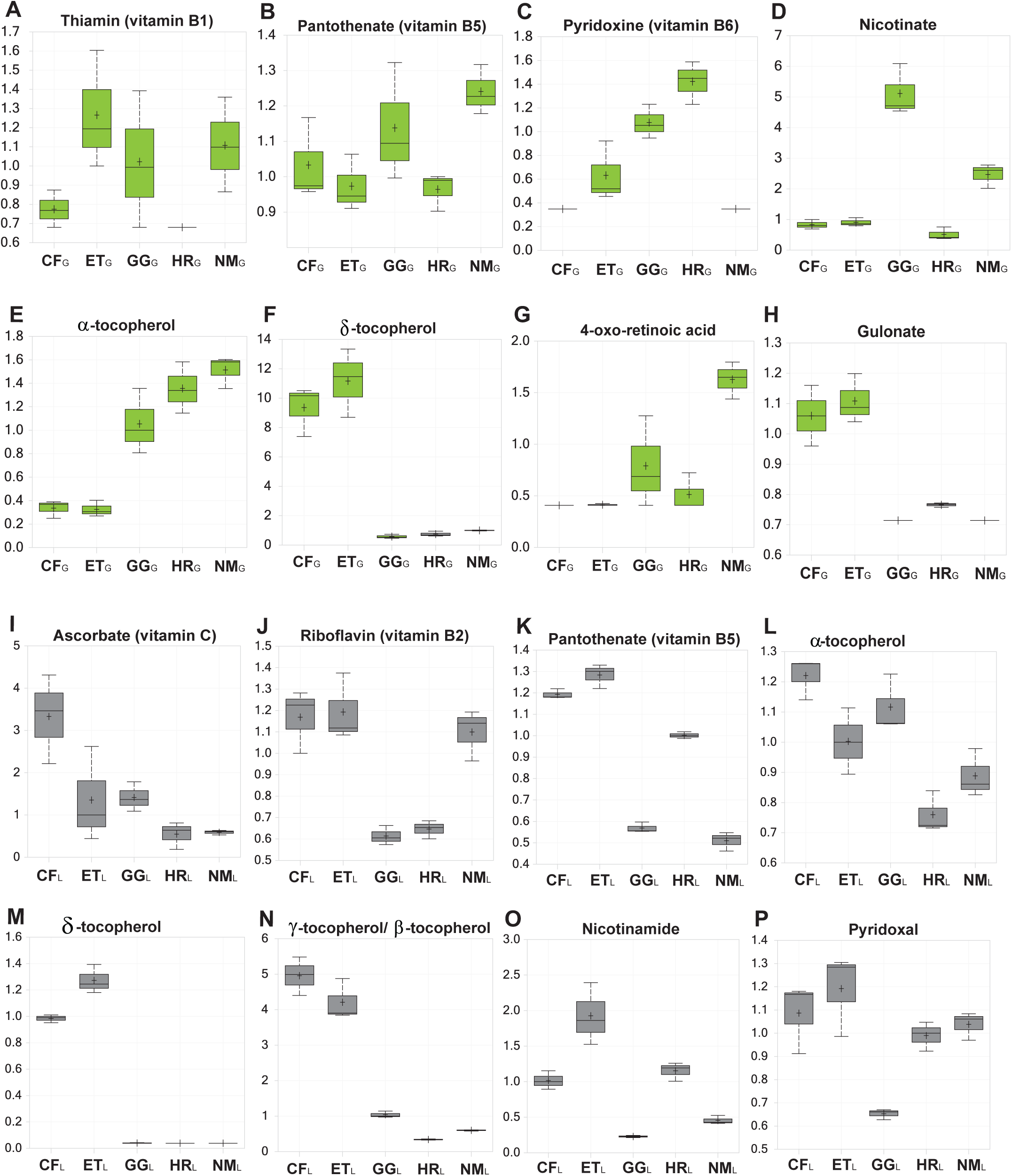
Contents of vitamins or vitamin-related metabolites detected in grains and leaves of the amaranth cultivars. A) Thiamin, B) pantothenate, C) pyridoxine, D) nicotinate, E) α-tocopherol, F) δ-tocopherol, G) 4-oxo-retinoic acid, and H) gulonate in grains. I) Ascorbate, J) riboflavin, K) pantothenate, L) α-tocopherol, M) δ-tocopherol, N) γ-tocopherol/β-tocopherol, O) nicotinamide, and P) pyridoxal in leaves. CF, Coral Fountain; ET, Emerald Tassels; GG, Golden Giant; HR, Hopi Red Dye; NM, New Mexico. (_G_), grain; (_L_), leaf.

### Secondary metabolites are detected in the amaranth grain and leaf tissues

Various secondary metabolites from plants have been associated with nutritive value, often reported to have antioxidant and anti-inflammatory properties, potentially leading to enhanced cardiovascular health, among other benefits [24]. Polyphenols, particularly from the flavonoid class, have been widely studied in this respect [25]. Among the five amaranth cultivars, we observed considerable variation in the contents of some flavonoid compounds, although we only detected five in this study (Fig. 4A-E). The level of flavonoids in amaranth grain was fairly low, as many flavonoids that are typically detected in grain crops were not detected at all in amaranth; even for those we detected, individual samples often had no data. We therefore suspect that the overall nutritional contribution of this compound class in amaranth grain may be minor at best. In amaranth leaf tissues, we observed considerable variation in flavonoid contents among cultivars; however, we only detected two compounds of the flavonoid class in amaranth leaves (Fig. 4F, G), both of which were highest in GG_L_.

**Fig. 4.**
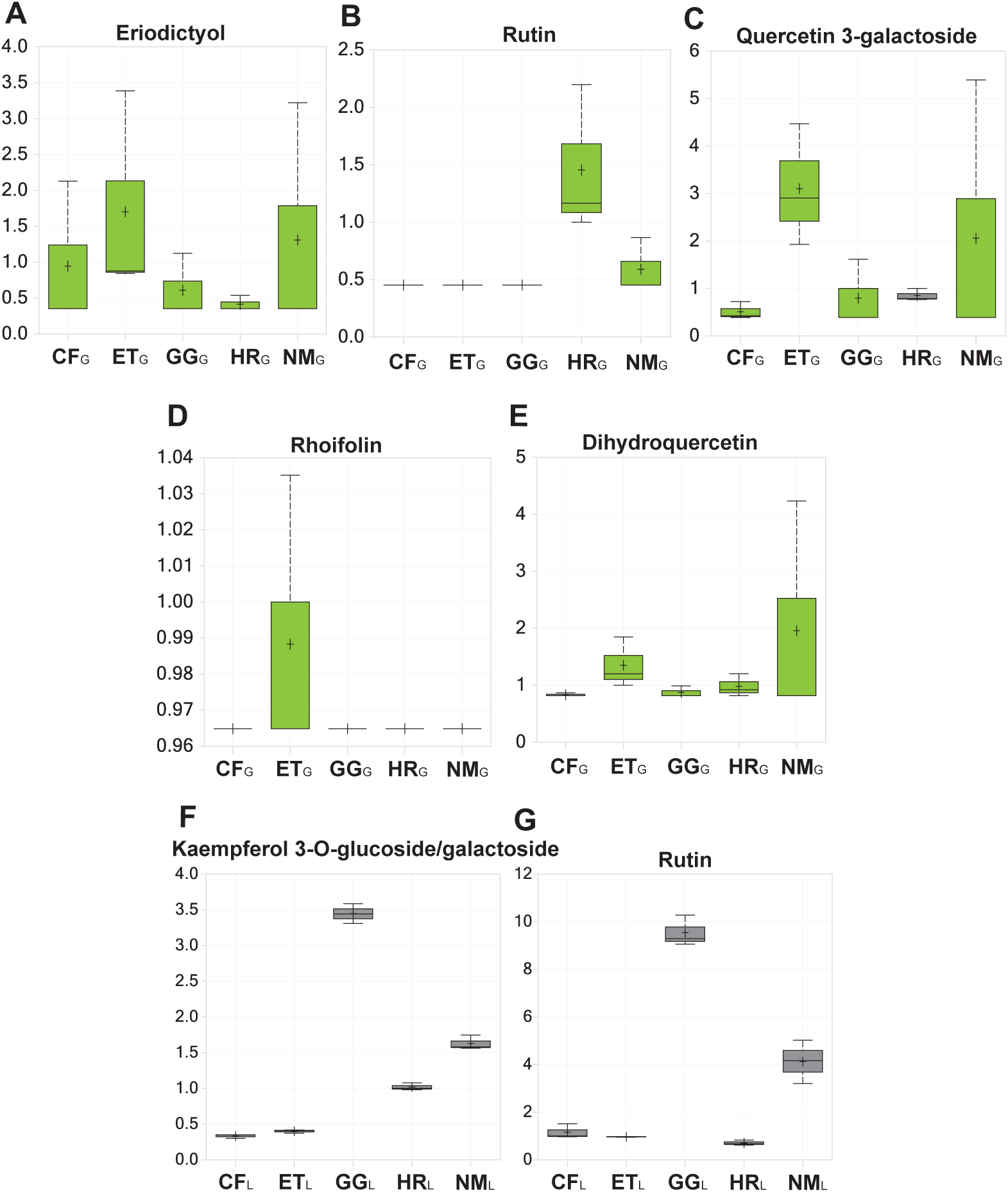
Contents of flavonoid-related compounds detected in grains and leaves of the amaranth cultivars. A) Eriodictyol, B) rutin, C) quercetin 3-galactoside, D) rhoifolin, and E) dihydroquercetin in grains. F) Kaempferol 3-O-glucoside/galactoside and G) rutin in leaves. CF, Coral Fountain; ET, Emerald Tassels; GG, Golden Giant; HR, Hopi Red Dye; NM, New Mexico. (_G_), grain; (_L_), leaf.

Benzenoids and chlorogenic acids (hydroxycinnamic acids esterified to quinate) were two classes of secondary metabolites that were widely represented in amaranth grain and leaf tissue and that displayed significant variations among cultivars. Some benzenoid catabolites (Fig. 5A, B) may contribute to the formation of volatile flavor compounds [26]; others have antioxidative properties useful for the preservation of lipids in foods [27]. Dietary salicylate (which protects plants from disease) was suggested to be beneficial to human heart health, much in the same way as low-dose aspirin [28]. The variation in the contents of these benzenoid catabolites among the grains of amaranth cultivars was significant, with salicylate and its glucoside as well as gentisic acid-5-glucoside being 10- to 30-fold higher in CF_G_ and ET_G_ relative to the other cultivars (Fig. 5C-F). However, the variation of these compounds in amaranth leaves was relatively minor, being typically around two-fold or less between any two cultivars (Fig. 5G–L).

**Fig. 5.**
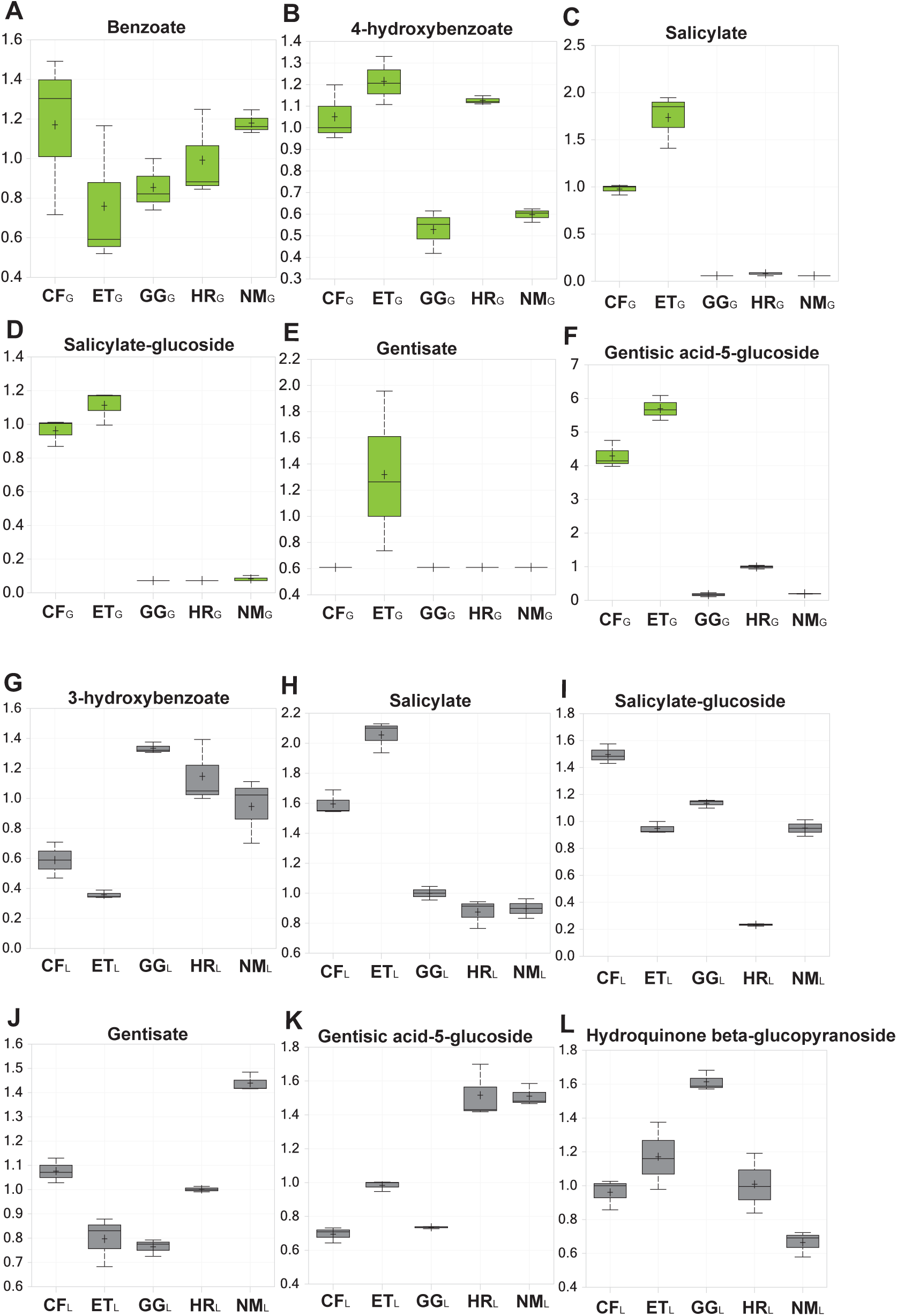
Contents of benzenoid metabolites detected in grains and leaves of the amaranth cultivars. A) Benzoate, B) 4-hydroxybenzoate, C) salicylate, D) salicylate-glucoside, E) gentisate, and F) gentisic acid-5-glucoside in grains. G) 3-hydroxybenzoate, H) salicylate, I) salicylate-glucoside, J) gentisate, K) gentisic acid-5-glucoside, and L) hydroquinone beta-glucopyranoside in leaves. CF, Coral Fountain; ET, Emerald Tassels; GG, Golden Giant; HR, Hopi Red Dye; NM, New Mexico. (_G_), grain; (_L_), leaf.

Chlorogenic acids are widely reported to have beneficial effects on cardiovascular, liver, and kidney health [29] as well as to promote cognitive function and neuroprotection [30]. Most of these studies [29–31] utilized the most abundant form found in coffee and chocolate, in which the class is named *trans*-5-O-caffeoyl-D-quinate (chlorogenic acid), but many chemical variants exist depending on the plant species in which they arise. We did not specifically detect chlorogenic acid in this study but instead measured a variety of isomers in which the hydroxycinnamic moiety was ferulate (Fig. 6) with an indication of the presence of 4-hydroxycinnamate (*p*-coumarate) forms as well. The various feruloylquinate isomers (up to five forms) could not be structurally distinguished in our system because of the absence of chemical standards; nonetheless, in almost all cases, the isomers contents were highly correlated to each other. The isomers may represent different chiral forms within the hydroxycinnamic acid moiety or, more likely, positional isomers for the site of esterification on quinate (or both). In Fig. 6, we show boxplots for the contents of the precursors (quinate, 4-hydroxycinnamate, ferulate) and for the esterified products feruloylquinate and coumaroylquinate in grain and leaf tissues. The phenylalanine ammonia-lyase (PAL) pathway provides *p*- coumarate for the formation of the hydroxycinnamic acids before their CoA-dependent esterification to quinate. The quinate content was much higher in the grain and leaf tissues of CF and ET, and the levels of the esterified products were associated with the quinate levels in all cases but not with the hydroxycinnamic acid levels. This finding indicates that the levels of feruloylquinate appeared to be driven by the quinate levels alone. This observation suggests that the regulation of this pathway relies on effects on the shikimate pathway upstream of phenylalanine ammonia-lyase (PAL).

**Fig. 6.**
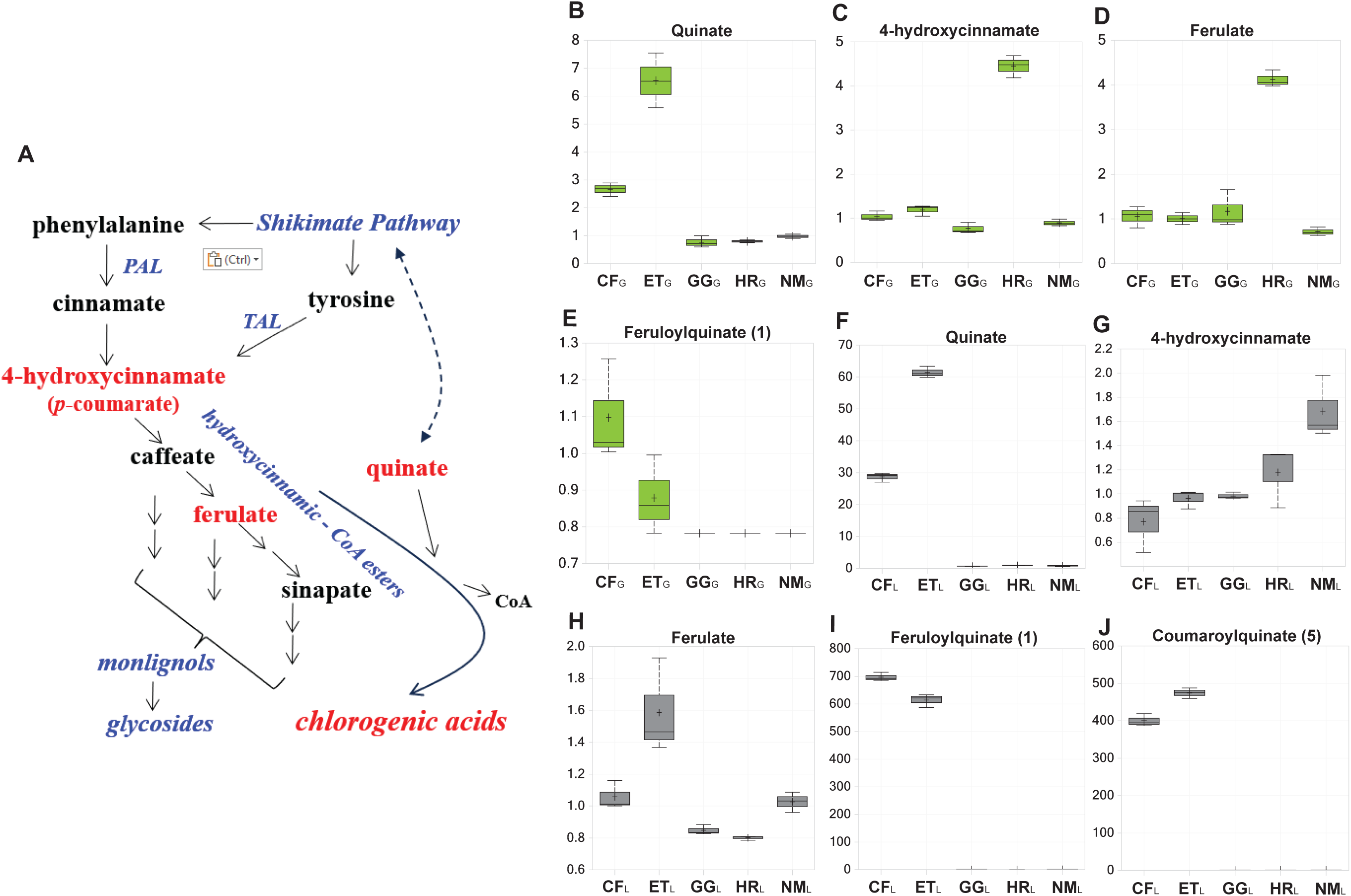
Contents of chlorogenic acids in the grains and leaves of amaranth cultivars. A) Diagram of the biochemical pathway leading to the formation of chlorogenic acids. B–J) Contents of compounds related to the biosynthesis of chlorogenic acids: B) quinate, C) 4-hydroxycinnamate, D) ferulate, and E) feruloylquinate (1) in grains. F) Quinate, G) 4-hydroxycinnamate, H) ferulate, I) feruloylquinate (1), and J) coumaroylquinate (5) in leaves. CF, Coral Fountain; ET, Emerald Tassels; GG, Golden Giant; HR, Hopi Red Dye; NM, New Mexico. (_G_), grain; (_L_), leaf.

### Anti-nutrients were detected in the amaranth grain and leaf tissues

Raffinose family oligosaccharides (RFOs) occur in many plants (see the biosynthetic pathway outline in Fig. S6), being especially ubiquitous in legumes, but their relatively high levels in amaranth grain is well known, and the fact that they are generally regarded as anti-nutrients may have some influence on the popularity of this grain as human food [32]. Looking at the relative levels of RFOs in the amaranth grain, the amount of raffinose was not strongly different between the cultivars, while CF_G_ had significantly more stachyose and verbascose than the four other cultivars; GG_G_ appeared to have the lowest levels of stachyose (Fig. S6). In amaranth leaves, we only detected stachyose, which was highest in HR_L_, as was its direct precursor galactinol, both of which were lowest in ET_L_ (Fig. S6). Another carbohydrate-related compound, oxalate (ethanedioate), is known to be high in amaranth grain. This compound has anti-nutritive properties and is considered toxic in high amounts, and routine ingestion of oxalate-rich foods can lead to depletion of important divalent cations, such as calcium, as well as the formation of kidney stones [33]. There was only minor variation in oxalate levels between the amaranth grain groups, with ET_G_ having the highest amount (Fig. S7). However, in the amaranth leaves, we detected considerable variation in oxalate levels among cultivars, with CF_L_ and HR_L_ having low amounts and NM_L_ having the highest amount (Fig. S7).

### Generation of a high-quality reference genome for the five amaranth cultivars

To generate high-quality reference genomes for the five cultivars, we extracted high-quality genomic DNA samples from fresh leaf tissues. We sequenced the five genomes and generated an average of 2.9 million PacBio HiFi circular consensus sequencing (CCS) reads with a median length of 18.9 kb, which corresponds to an average of 53.44 Gb of raw sequence data per cultivar. Assuming a genome size of 466 Mb for *A. hypochondriacus* [34], the corresponding expected genome coverage would be ∼115 × (Table 2). The primary genome assembly consisted of 1,064–1,667 contigs with an N50 range of 2.3–3.9 Mb (Table 2) and a total size between 465 Mb and 483 Mb.

**Table 2.** Summary statistics of the five amaranth genome assemblies.

| **Assembly statistics** | **CF** | **ET** | **GG** | **HR** | **NM** |
| --- | --- | --- | --- | --- | --- |
| # HiFi CCS reads | 2,956,722 | 2,604,135 | 2,626,199 | 3,630,959 | 2,735,384 |
| HiFi CCS reads (Gb) | 57.9 | 47.5 | 52.6 | 69.5 | 39.7 |
| Coverage | 124 × | 101 × | 112 × | 149 × | 85 × |
| Total number of contigs | 1093 | 1064 | 1130 | 1195 | 1667 |
| Total length (bp) | 468,059,521 | 465,071,770 | 483,580,607 | 480,005,045 | 476,067,050 |
| # contigs (≥ 25,000 bp) | 1078 | 1055 | 1120 | 1183 | 1447 |
| Total length (≥ 25,000 bp) | 467,723,944 | 464,862,660 | 483,356,820 | 479,730,140 | 471,160,992 |
| # contigs (≥ 50,000 bp) | 713 | 661 | 766 | 685 | 420 |
| Total length (≥ 50,000 bp) | 452,912,348 | 449,349,769 | 469,058,912 | 459,758,129 | 436,398,612 |
| Largest contig (bp) | 14,025,610 | 12,975,424 | 15,060,313 | 15,677,540 | 18,573,590 |
| GC (%) | 34.0 | 34.2 | 34.0 | 33.3 | 33.2 |
| N50 | 3,057,888 | 2,328,578 | 2,554,507 | 3,899,886 | 3,368,805 |
| N90 | 174,614 | 172,550 | 153,969 | 113,063 | 97,185 |
| auN | 5,411,356.8 | 5,396,009.2 | 4,808,603.4 | 6,155,777.9 | 5,894,610.2 |
| L50 | 28 | 30 | 37 | 25 | 27 |
| L90 | 308 | 311 | 343 | 267 | 294 |
| # Ns per 100 kb | 0 | 0 | 0 | 0 | 0 |

### Structural variation (SV) and transposable element (TE) analysis of the amaranth genomes

We used the newly assembled NM genome as a reference for comparison to the four other genome assemblies to identify insertions, deletions, and inversions. The number of genomic regions not shared between each of the four cultivars and NM ranged from 7,478 (GG) to 53,411 (CF), likely reflecting the relative evolutionary distance between each cultivar and the reference genome (Table S8). Cultivars CF and ET appeared to be distantly related to NM, whereas GG was more closely related to NM, as we detected fewer variants between them (Table 1).

We then investigated the distance between SVs and 5’ end of the nearby highly supported (posterior probability greater than 0.4) genes. In the case of the distantly related CF and ET species, 26,8 % and 33.8 % of their genes have an insertion or a deletion at less of 3 Kb from the gene 5’ end. For GG and HR these percentages dropped to 6.7% and 12.8%, respectively reflecting the fewer number of SVs detected in those cultivars when compared to NM. We screened the five assembled genomes for TEs, as they are considered important components of plant genomes. Overall, the TE content of the five genome assemblies ranged from 46.9% to 48.7%, with long terminal repeat (LTR)-type *Copia* (8.6–10.3%) being the most abundant TE class in all cultivar genomes. Most of the TE classes showed slight variation in abundance among the five cultivars (Table S9).

### De novo prediction of protein-coding genes

Iso-seq reads generated from the leaf, flower, and root tissues of the five cultivars (Table S10) were used as extrinsic data to support the prediction of protein-coding genes in the five genomes using Augustus [44], included within OmicsBox 2.2.4 [28]. Overall, we identified 40,472 - 72,088 putative genes (Table S4). Of these 19,921- 50,606 were functionally annotated (Table S11), and were assigned to the corresponding gene ontology (GO) terms (Table S12).

## Discussion

The nutritional value of amaranth varies widely depending on a range of factors, including the genetic make-up, growth environment, agronomic inputs, post-harvest handling, and processing. These factors affect the levels of protein, carbohydrates, fats, minerals, vitamins, fiber, and bioactive compounds such as antioxidants in the edible parts (grain and leaf tissues). The untargeted metabolomic profiling conducted in this study revealed significant differences in grain and leaf metabolites among the five amaranth cultivars, providing valuable insights into their nutritional quality and potential health-promoting properties. We identified 495 compounds in grains and 559 in leaves, encompassing a wide range of metabolic pathways crucial for plant growth and development, and with benefits to human health. Pairwise comparisons revealed significant variations in metabolite composition, with distinct patterns observed among the cultivars. A principal component analysis corroborated these findings, highlighting clear distinctions between cultivars based on their metabolic contents. Notably, cultivars CF and ET displayed higher overall metabolite contents compared to the other cultivars across various compound classes, suggesting potential superiority in their nutritional composition and physiological properties.

Our analysis also revealed cultivar-related variations in key pathways associated with nutrition and stress mitigation. For instance, we observed differences in the contents of sulfur-containing compounds, such as methionine and its derivatives, among the cultivars, indicating potential variations in protein quality and sulfur metabolism. Furthermore, variations in the contents of stress-related metabolites, including glutathione and quaternary ammonium compounds, may have agronomic implications, highlighting cultivar-specific adaptations to environmental stressors such as drought. Assessment of vitamin contents showed variation among cultivars, with no clear pattern suggesting the superiority of one cultivar over others. However, we did observe cultivar-specific differences in the levels of vitamins B1, B5, B6, alpha-tocopherol, delta-tocopherol, and other vitamin-related metabolites in either grain or leaf tissues. Similarly, secondary metabolites, such as flavonoids, benzenoids, and chlorogenic acids, exhibited significant variation among cultivars, indicating potential differences in their antioxidant and anti-inflammatory properties.

Analysis of anti-nutrient contents revealed variation in the levels of RFOs and oxalate among cultivars, with CF and ET displaying distinct profiles compared to the other cultivars. These findings shed light on the potential health implications of consuming different amaranth cultivars and underscore the importance of selecting cultivars with favorable nutritional profiles for human consumption. Overall, this study provides valuable insights into the metabolic diversity of amaranth cultivars and highlights the potential of metabolomic approaches for assessing the nutritional quality and health-promoting properties of alternative crops like amaranth. Future research should focus on elucidating the genetic and environmental factors underlying the observed variations and their implications for crop improvement and food security. Amaranth possesses abundant metabolites with health-promoting properties; however, it is essential to manage anti-nutritional factors present in amaranth. Clustered regularly interspaced short palindromic repeats (CRISPR)/CRISPR-associated nuclease (Cas)-based technologies offer the potential to decrease or abolish the accumulation of anti-nutrients by knocking out or lowering the expression of their biosynthetic genes. Therefore, in this work, we sequenced and assembled a high-quality reference genome for these five amaranth cultivars that were not previously available. We analyzed the repetitive sequences in these genomes and their sequence variation, which provides valuable insights into the complexity and diversity of those cultivars. These data are essential for future genomic studies and for understanding the molecular mechanisms underlying traits of interest and various biological pathways. In addition, these data will facilitate the identification of the key genes controlling different agronomic and nutritional attributes of amaranth, which enables the application of gene editing technologies to improve those cultivars, thereby paving the way for establishing them as superfood crops.

## Conclusion

Amaranth has recently gained attention as a promising crop to enhance global food security due to its remarkable nutritional profile and adaptability to adverse environmental conditions. With its richness in essential amino acids, proteins, dietary minerals, and bioactive compounds, amaranth offers a valuable solution for addressing nutritional deficiencies and dietary restrictions such as gluten sensitivity. Moreover, its edible grains and leaves enhance its versatility and potential as a sustainable food source. By employing metabolomics and whole-genome sequencing techniques, this study fills this knowledge gap and identifies cultivars with the highest content of health-promoting compounds. Such insights are crucial for targeted trait engineering and the development of cultivars optimized for nutritional value and agronomic performance, thereby reaching food security and sustainability in the face of increasing agricultural challenges.

## Materials and Methods

### Plant materials and extraction of sample metabolites

Five grain amaranth cultivars were employed in this study: *Amaranthus caudatus* ‘Coral Fountain’ (CF) and ‘Emerald Tassels’ (ET); *A. cruentus* ‘Golden Giant’ (GG) and ‘Hopi Red Dye’ (HR); and *A. hypochondriacus* ‘New Mexico’ (NM). Mature grains were obtained from Hudson Valley Seed Company, United States, and directly used for the metabolome profiling. Grains were also sown and plants cultivated at the greenhouse of the King Abdullah University of Science and Technology, Saudi Arabia for analyzing the metabolome content of leaves. Mature grains and leaves of two-month-old plants were collected from nine individual plants for each cultivar. The samples from three individuals were pooled as one biological replicate and three replicates were analyzed. Grains and leaves were ground into fine powder in liquid nitrogen and then lyophilized for 30 h using FreeZone 2.5 Plus (Labconco). Samples were then prepared by Metabolon using the automated MicroLab STAR system from Hamilton Company. Several recovery standards were added prior to extraction as a means of quality control. Samples were subjected to methanol extraction with vigorous shaking for 2 min using GenoGrinder 2000 (Glen Mills Inc) to precipitate protein and dissociate small molecules bound to protein, followed by centrifugation. The resulting extract was divided into five fractions: two for analysis by two separate reverse-phase (RP)/UPLC–MS/MS methods using positive ion mode electrospray ionization (ESI), one for analysis by RP/UPLC–MS/MS using negative ion mode ESI, one for analysis by HILIC/UPLC–MS/MS using negative ion mode ESI, and one reserved for backup. Samples were placed briefly on a TurboVap (Zymark) to remove the organic solvent. The extracts were stored overnight under nitrogen before preparation for analysis.

### Ultra-high performance liquid chromatography-tandem mass spectrometry (UPLC-MS/MS)

All procedures used a Waters ACQUITY ultra-performance liquid chromatography (UPLC) system coupled to a Thermo Scientific Q-Exactive high-resolution/accurate mass spectrometer interfaced with a heated electrospray ionization (HESI-II) source and an Orbitrap mass analyzer operated at a resolution of 35,000 mass units. The extracted sample was dried and then reconstituted in solvents compatible with each of the four methods. Each reconstitution solvent incorporated a set of standards at consistent concentrations to ensure uniform injection and chromatographic performance. One aliquot of the sample was subjected to analysis under acidic positive ion conditions, optimized for hydrophilic compounds. Here, the extract was eluted in a gradient manner from a C18 column (Waters UPLC BEH C18-2.1 × 100 mm, 1.7 µm) using a mixture of water and methanol containing 0.05% (v/v) perfluoropentanoic acid (PFPA) and 0.1% (v/v) formic acid (FA). The second aliquot of the sample was also analyzed using acidic positive ion conditions but was specifically optimized for more hydrophobic compounds. In this scenario, the extract was eluted from the same C18 column using a mixture of methanol, acetonitrile, water, 0.05% (v/v) PFPA, and 0.01% (v/v) FA, with an overall higher organic content. The third aliquot was analyzed using basic negative ion mode optimized conditions using a separate dedicated C18 column. The basic extracts were eluted from the column employing methanol and water containing 6.5 mM ammonium bicarbonate at pH 8. The fourth aliquot was analyzed via negative ionization mode following elution from a HILIC column (Waters UPLC BEH Amide 2.1 × 150 mm, 1.7 µm) using a gradient consisting of water and acetonitrile with 10 mM ammonium formate and at a pH of 10.8. The mass spectrometry analysis alternated between MS and data-dependent MS^n^ scans using dynamic exclusion. The scan range varied slightly between methods, but covered approximately 70–1000 *m/z*.

### Data extraction and compound identification

Raw data were extracted, peak-identified, and processed for quality control using Metabolon hardware and software suite. Compounds were identified by comparing them to a library comprising over 4,500 commercially available purified standard compounds. Furthermore, additional entries in the mass spectral database were generated for structurally unidentified biochemicals, recognized by their recurring chromatographic and mass spectral characteristics. Various curation procedures were implemented to ensure the assembly of a high-quality dataset suitable for statistical analysis and data interpretation, encompassing (1) rigorous validation of precise and consistent identification of genuine chemical entities and (2) elimination of system-induced artifacts, erroneous assignments, redundancies, and background noise. Quantification of peaks was conducted based on the area-under-the-curve detector ion counts.

### Statistical analysis of amaranth metabolites

The data were analyzed using Welch’s two-sample *t*-tests in GraphPad Prism 9.3.1 software. A *P-*value of <0.05 was used as threshold for statistical significance. In cases of missing values, if any, imputation was performed using the observed minimum value for the respective compound. Principal component analysis (PCA) was employed for data exploration. Scaled imputed data for all 15 amaranth grain and 15 leaf samples were plotted, with the compounds arranged by general metabolic category (Super Pathway). In the heatmap, the median value across all samples was rendered in white, with higher values represented by varying shades of red (up to saturation at 3-fold) and lower values indicated by gradations of blue (with 0.33 of the median serving as saturation). We chose to use the median (rather than the overall mean) because it more realistically represents the general metabolite yield without being biased by a few compounds with very extreme differences.

### PacBio library construction, genome sequencing, and assembly

The seeds of all five cultivars were planted in soil under standard greenhouse conditions at the greenhouse facility of the King Abdullah University of Science and Technology, Saudi Arabia. The resulting seedlings were grown for six weeks. Approximately 20 g of tissue from young leaves was collected from each cultivar following a 48-h dark treatment to remove starch and immediately flash frozen in liquid nitrogen and then stored at −80°C until DNA extraction. High-molecular-weight genomic DNA was isolated using a modified cetyltrimethylammonium bromide (CTAB) method as previously described [15, 21]. The size of the extracted DNA and susceptibility to restriction enzyme digestion were evaluated by pulsed-field electrophoresis (CHEF) on 1% (w/v) agarose gels; the DNA concentration was determined using a Qubit fluorometer (Thermo Fisher Scientific, Waltham, MA). The high-molecular-weight DNA was fragmented into large fragments (10–30 kb) using Covaris g-TUBE according to the manufacturer’s instructions (Pacific Biosciences) and subsequently purified with PB Beads (PacBio, Menlo Park, CA). Sequencing libraries were constructed following the manufacturer’s protocol using a SMRTbell Express Template Prep kit 2.0. The libraries were quantified using a Qubit dsDNA high-sensitivity assay (Invitrogen, USA), and size verification was conducted on a Femto Pulse System (Agilent). Genomic DNA from all five cultivars was sequenced on a PacBio Sequel II system using SMRT Cell 1M chemistry version 3.0 in CCS mode for a total of 30 h at the Core lab of the King Abdullah University of Science and Technology. Raw data generated ranges from 39.7 to 69.5 corresponding to coverage ranging from 85X to 149X (Table 2). The five amaranth genomes were *de novo* assembled using Hifiasm v. 0.15.5 run under default settings. The most important assembly metrics were evaluated by running Quast [35] and carrying out the BUSCO analyses [36] using the embryophyta_odb10 BUSCO gene set.

### Structural variation (SVs) and transposable element (TE) quantification and masking

Structural variation analysis was performed using the assembled amaranth NM genome as a reference, and the presence and absence of structural variants were analyzed using NGMLR [37] as an aligner and SVIM [38] as a caller. TE libraries were created for each cultivar using the tool EDTA [39]. TE libraries (with *Helitron* predictions removed) were then used to assess the TE content in the five genomes using RepeatMasker [40]. The soft-masked genome assemblies were then used for gene prediction.

### Iso-seq library preparation and sequencing for baseline genome annotation

Total RNA was extracted from the leaf, flower, and root tissues of plants for the five cultivars using a Maxwell RSC Plant RNA Kit and a Maxwell RSC48 instrument following Promega’s protocol. The PacBio Iso-seq sequencing library was prepared following the standard protocol for isoform sequencing using the SMRTbell Express Template Prep Kit 2.0. cDNA samples from the three tissues were then barcoded and amplified. The multiplexed library was prepared for sequencing using a PacBio Sequel II Sequencing kit 2.0, loaded onto six 1M SMRT cells, and sequenced in CCS mode in a PacBio Sequel II instrument for 30 h. The Iso-seq raw reads were filtered and clustered into high-quality (HQ), non-chimeric, full-length transcripts using the isoseq3 v.3 pipeline [26].

### De novo gene-coding annotation with long and short RNA reads

For *de novo* gene prediction, the Gene Finding module was run using the genome assembly soft-masked for TEs and Iso-seq as direct evidence using Augustus [27] with default settings as implemented in OmicsBox 2.2.4 [28]. The translated annotation was assessed using BUSCO v.3.0 [29] against the Viridiplantae database v.10. the functional annotation of the predicted genes followed by gene ontology (GO) analysis was performed using InterProScan version 5.39 with default settings, and all the information was integrated using the OmicsBox genome analysis suite to generate the corresponding reports.

## Declarations

### Ethics approval and consent to participate

Not applicable.

### Consent for publication

Not applicable.

### Availability of data and materials

The MS metabolomic datasets generated during the current study are available in the MetaboLights database with the identifier numbers MTBLS815 and MTBLS3498. The genome assembly of the five amaranth varieties and the Isoseq reads are available in the NCBI; bioproject PRJNA1226974.

## Authors’ contributions

MM and KS conceived the project; KS, AZ, UT, KrS, and SK conducted the experiments; KS, AZ, and LR analyzed the data; and KS, MM, and AZ wrote the manuscript; KS prepared the figures; KS, MM, AZ, and RW reviewed the manuscript.

## Acknowledgments

We thank Nahed Mohamed and the Laboratory for Genome Engineering and Synthetic Biology at KAUST for critical discussions and technical help with this work. We thank Hudson Valley Seed Company, United States, for providing the seeds and panicle images.

