## Supplementary material for "Nutritional quality and genetic differences of five amaranth cultivars revealed by metabolome profiling and whole-genome sequencing": Fig S2

**PCA component 1 main drivers**

| SUPER_PATHWAY | CHEMICAL_NAME | PC1 |
|---|---|---|
| Amino Acid | **quinate** | 0.06607 |
| Amino Acid | **5-methylthioribose**** | 0.06599 |
| Cofactors | **dehydroascorbate** | 0.06559 |
| Cofactors | **delta-tocopherol** | 0.06646 |
| Secondary metabolism | **coumaroylquinate (2)** | 0.06663 |
| Secondary metabolism | **coumaroylquinate (5)** | 0.0667 |
| Secondary metabolism | **coumaroylquinate (3)** | 0.06648 |
| Secondary metabolism | **feruloylquinate (1)** | 0.06578 |
| Secondary metabolism | **feruloylquinate (3)** | 0.06647 |
| Secondary metabolism | **feruloylquinate (4)** | 0.06588 |

| SUPER_PATHWAY | CHEMICAL_NAME | PC1 |
|---|---|---|
| Amino Acid | **glycine** | -0.06121 |
| Amino Acid | **shikimate** | -0.06262 |
| Amino Acid | **o-tyramine** | -0.0642 |
| Amino Acid | **isoleucine** | -0.05895 |
| Carbohydrate | **galactinol** | -0.0595 |
| Carbohydrate | **glucarate (saccharate)** | -0.0622 |
| Nucleotide | **uridine 5'-monophosphate (UMP)** | -0.05954 |
| Peptide | **prolylalanine** | -0.06094 |
| Secondary metabolism | **3-hydroxybenzoate** | -0.06067 |
| Secondary metabolism | **kaempferol 3-O-glucoside/galactoside** | -0.06136 |

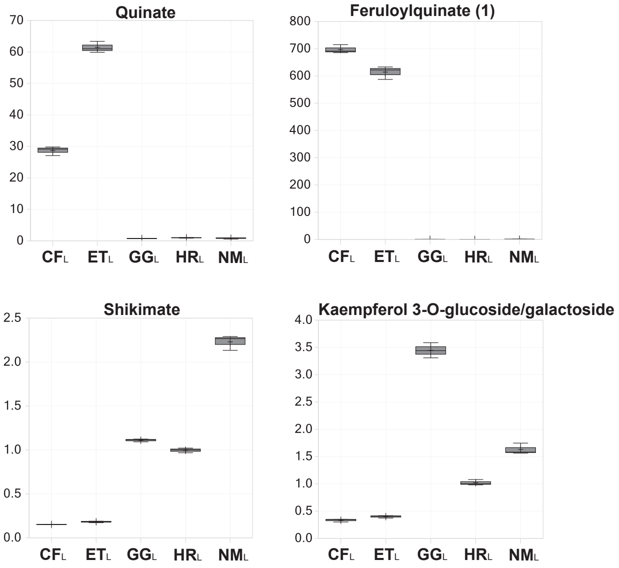


**PCA component 2 main drivers**

| SUPER_PATHWAY | CHEMICAL_NAME | PC2 |
|---|---|---|
| Amino Acid | **N-acetylglutamate** | 0.082733 |
| Amino Acid | **ornithine** | 0.081247 |
| Lipids | **1-stearoyl-2-oleoyl-GPC (18:0/18:1)** | 0.083412 |
| Lipids | **1-oleoyl-2-linoleoyl-GPC (18:1/18:2)*** | 0.081927 |
| Lipids | **1,2-dilinoleoyl-GPE (18:2/18:2)*** | 0.086101 |
| Lipids | **1-palmitoleoyl-2-linolenoyl-GPG (16:1/18:3)*** | 0.083106 |
| Lipids | **1,2-dilinoleoyl-galactosylglycerol (18:2/18:2)*** | 0.084997 |
| Cofactors | **nicotinamide ribonucleotide (NMN)** | 0.082082 |
| Secondary metabolism | **nicotianamine** | 0.082575 |
| Secondary metabolism | **beta-cryptoxanthin** | 0.084299 |

| SUPER_PATHWAY | CHEMICAL_NAME | PC2 |
|---|---|---|
| Amino Acid | **tryptophan** | -0.07069 |
| Amino Acid | **tyramine** | -0.07057 |
| Amino Acid | **homoserine lactone** | -0.06559 |
| Amino Acid | **saccharopine** | -0.07528 |
| Amino Acid | **S-methylmethionine** | -0.07138 |
| Amino Acid | **N6-acetyllysine** | -0.07771 |
| Amino Acid | **histidine** | -0.08707 |
| Carbohydrate | **stachyose** | -0.07867 |
| Lipids | **1-palmitoyl-GPI (16:0)** | -0.0658 |
| Secondary metabolism | **salidroside** | -0.07235 |

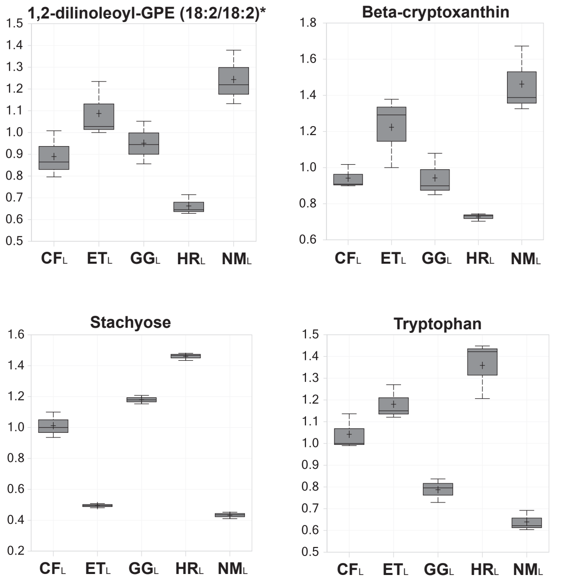


**Fig. S2. Compounds influencing separation along component 1 and component 2 of the leaf PCA plot.** The compounds with the top 10 positive and top 10 negative scores are listed.
