## Supplementary material for "Nutritional quality and genetic differences of five amaranth cultivars revealed by metabolome profiling and whole-genome sequencing": Fig S3

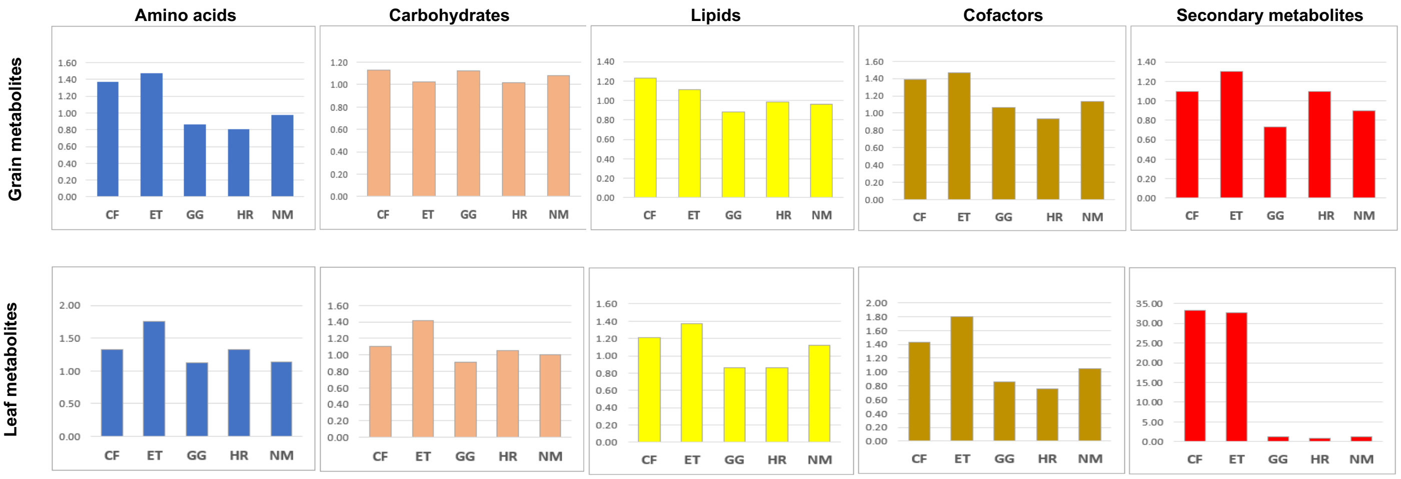


**Fig. S3. Average response by cultivar of all compounds classified in the five major pathway families (amino acids, carbohydrates, lipids, cofactors, secondary metabolites).**
