## Supplementary material for "Nutritional quality and genetic differences of five amaranth cultivars revealed by metabolome profiling and whole-genome sequencing": Fig S4

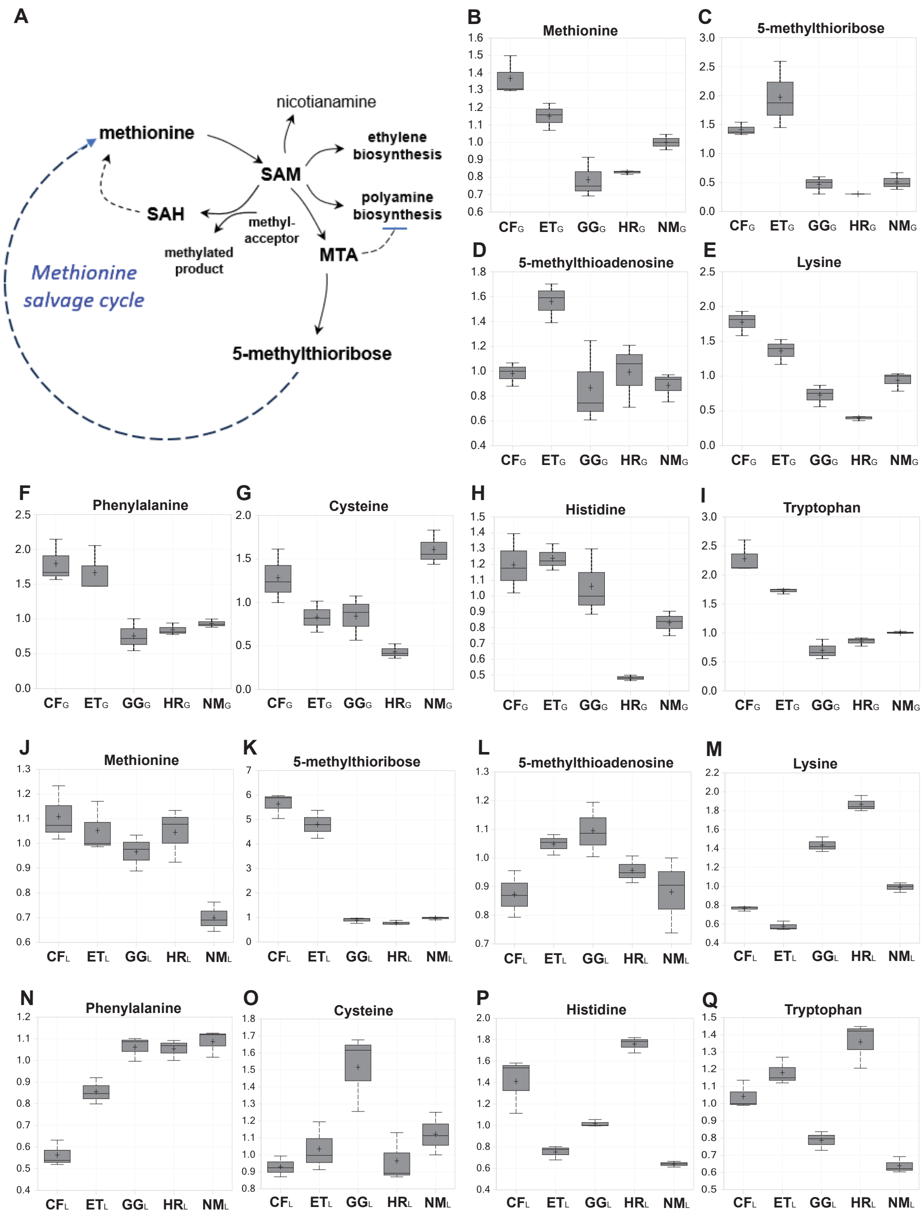


**Fig. S4. Representative amino acid and related compounds of the grain and leaf metabolites of the five amaranth cultivars.** A) Diagram of the methionine salvage cycle and related pathways. B–I) Contents in amino acids and derivatives of the grain metabolites. J–Q) Contents in amino acids and derivatives of the leaf metabolites.
