## Supplementary material for "Nutritional quality and genetic differences of five amaranth cultivars revealed by metabolome profiling and whole-genome sequencing": Fig S5

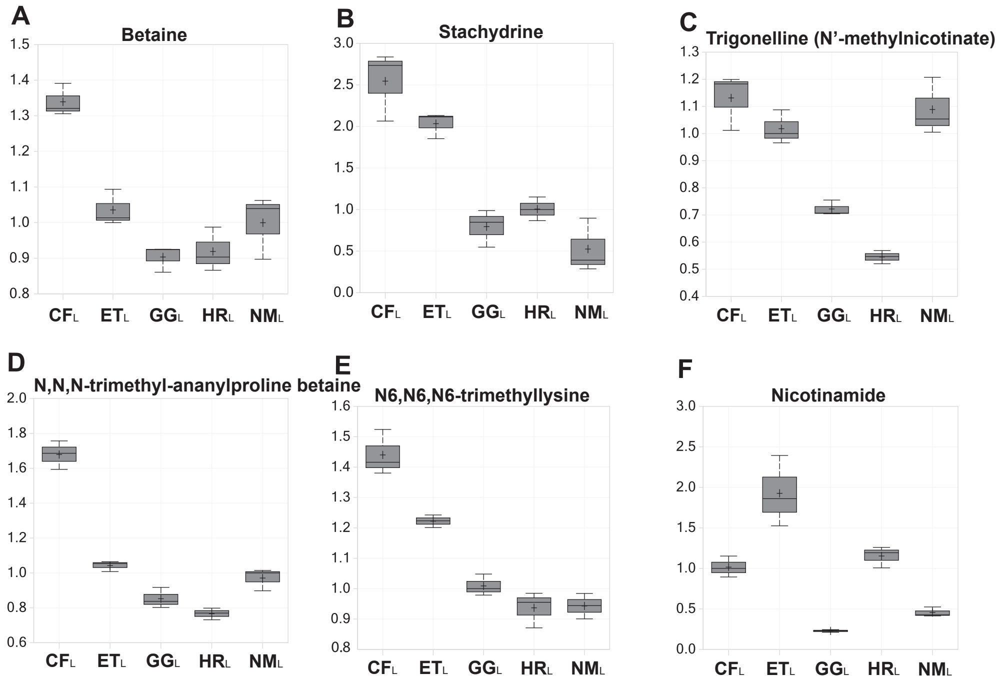


**Fig. S5.** Box plots for compounds associated with stress mitigation in the amaranth leaf of the five cultivars. A) betaine, B) stachydrine, C) trigonelline, D) N,N,N-trimethyl-ananylproline betaine, E) N6,N6,N6-trimethyllysine, F) nicotinamide.
