## Supplementary material for "Nutritional quality and genetic differences of five amaranth cultivars revealed by metabolome profiling and whole-genome sequencing": Fig S6

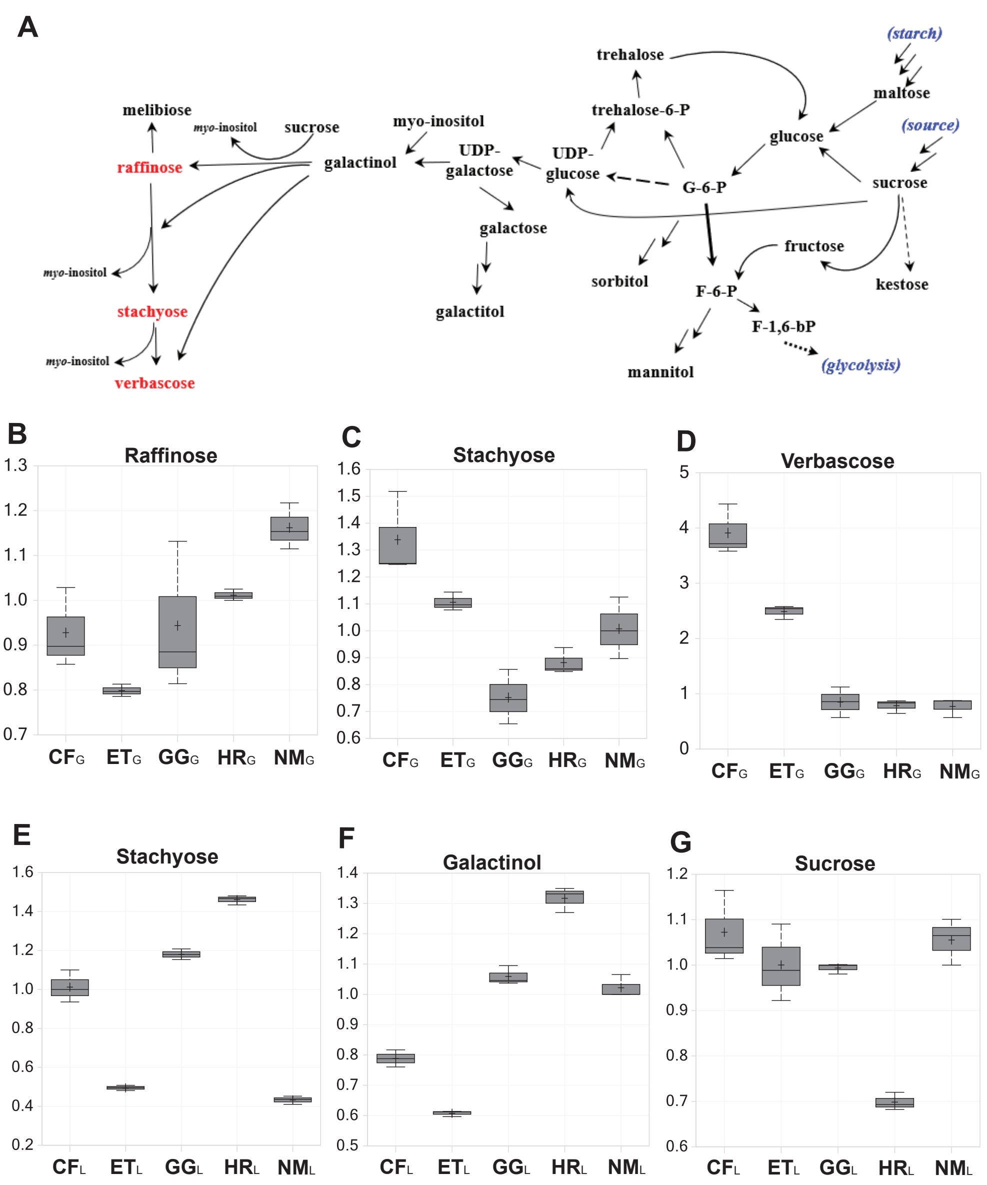


**Fig. S6. Carbohydrate metabolism and representative carbohydrate metabolites in Amaranth grain and leaf.** A) Simplified diagram of carbohydrate metabolism in plants, including the biosynthesis of raffinose family oligosaccharides (RFOs). B–D) Boxplot of the RFO contents in the amaranth grain samples. E–G) Boxplot of the RFO contents in the amaranth leaf samples.
