## Supplementary figures and images for "Nutritional quality and genetic differences of five amaranth cultivars revealed by metabolome profiling and whole-genome sequencing"

### Fig S7

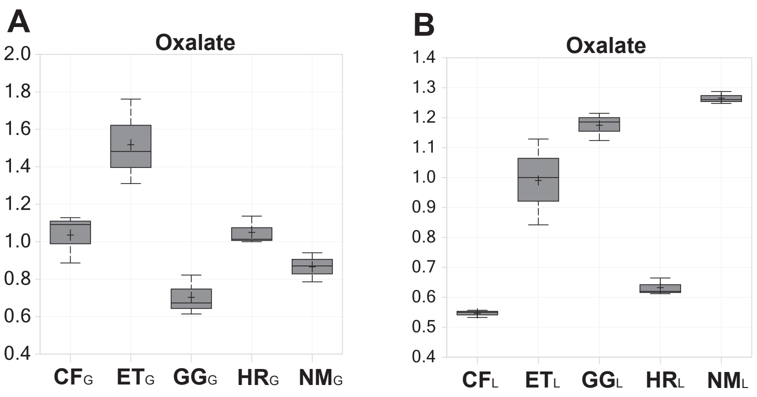


**Fig. S7.** Boxplot of the oxalate contents in the amaranth grain (A) or leaf (B).
