## Supplementary material for "Nutritional quality and genetic differences of five amaranth cultivars revealed by metabolome profiling and whole-genome sequencing": Table S7

**Table S1.** Ranking of vitamins or vitamin-related metabolites detected in the leaves of the five amaranth varieties. Compounds were scored based on the order of abundance (1=highest, 5=lowest). Below the table is the average across the compounds.

| Biochemical Name | CF | ET | GG | HR | NM |
|---|---|---|---|---|---|
| pantothenate | 2 | 1 | 4 | 3 | 5 |
| nicotinamide | 3 | 1 | 5 | 2 | 4 |
| NAD+ | 5 | 3 | 2 | 4 | 1 |
| riboflavin (vitamin B2) | 2 | 1 | 5 | 4 | 3 |
| ascorbate (vitamin C) | 1 | 3 | 2 | 5 | 4 |
| dehydroascorbate | 2 | 1 | 5 | 4 | 3 |
| gulonate* | 2 | 1 | 3 | 4 | 5 |
| alpha-tocopherol | 1 | 3 | 2 | 5 | 4 |
| delta-tocopherol | 2 | 1 | 3 | 4 | 4 |
| gamma-tocopherol/beta-tocopherol | 1 | 2 | 3 | 5 | 4 |
| pyridoxal | 2 | 1 | 5 | 4 | 3 |
| pyridoxamine | 4 | 1 | 5 | 3 | 2 |
| pyridoxamine phosphate | 2 | 1 | 4 | 5 | 3 |
| *av rank* | *2.23* | *1.54* | *3.69* | *4.00* | *3.46* |
