## Supplementary material for "Nutritional quality and genetic differences of five amaranth cultivars revealed by metabolome profiling and whole-genome sequencing": Table S8

**Table S2.** Number of insertions, deletions, and inversions between the four genomes and the New Mexico (NM) genome.

| Structure variant type | CF | ET | GG | HR |
|---|---|---|---|---|
| **Insertion** | 25,401 | 25,258 | 3,754 | 10,761 |
| **Deletion** | 27,973 | 27,910 | 3,720 | 11,857 |
| **Inversion** | 37 | 35 | 4 | 16 |
| **Total variants** | 53,411 | 53,203 | 7,478 | 22,634 |
