## Supplementary material for "Nutritional quality and genetic differences of five amaranth cultivars revealed by metabolome profiling and whole-genome sequencing": Table S10

**Table S10.** Summary statistics of the five amaranth Isoseq analysis

| Statistics | CF | ET | GG | HR | NM |
|---|---|---|---|---|---|
| **Isoseq reads** | 3,585,920 | 3,509,350 | 3,789,538 | 3,377,672 | 4,172,578 |
| **Isoseq reads (Gbp)** | 7,0 | 6,1 | 7,9 | 7,1 | 8,8 |
| **N50** | 2,157 | 2,059 | 2,312 | 2,374 | 2,383 |
