## Supplementary material for "Nutritional quality and genetic differences of five amaranth cultivars revealed by metabolome profiling and whole-genome sequencing": Table S11

**Table S11.** Number of predicted genes in the five amaranth genomes

| Statistics | CF | ET | GG | HR | NM |
|---|---|---|---|---|---|
| **Predicted genes** | 41,503 | 72,088 | 51,739 | 40,472 | 41,881 |
| **Predicted genes (pp > 0.4)** | 25,066 | 48,731 | 30,864 | 25,272 | 13,578 |
| **Predicted genes with functional annotation** | 19,921 | 50,606 | 31,106 | 19,921 | 21,132 |
