## Supplementary material for "Nutritional quality and genetic differences of five amaranth cultivars revealed by metabolome profiling and whole-genome sequencing": Fig S2

**PCA component 1 main drivers**

| SUPER_PATHWAY | CHEMICAL_NAME | PC1 |
|---|---|---|
| Amino Acid | methyl indole-3-acetate | 0.051716 |
| Amino Acid | 1-methyl-5-imidazolelactate | 0.067354 |
| Carbohydrate | alpha-ketoglutarate | 0.056013 |
| Carbohydrate | ribonate | 0.053787 |
| Carbohydrate | galactinol | 0.062513 |
| Lipids | 1-palmitoyl-2-stearoyl-GPC (16:0/18:0) | 0.050382 |
| Lipids | 1-stearoyl-2-oleoyl-GPI (18:0/18:1)* | 0.059053 |
| Cofactors | alpha-tocopherol | 0.062458 |
| Nucleotide | adenosine | 0.050947 |
| Nucleotide | N6-methyladenosine | 0.06456 |

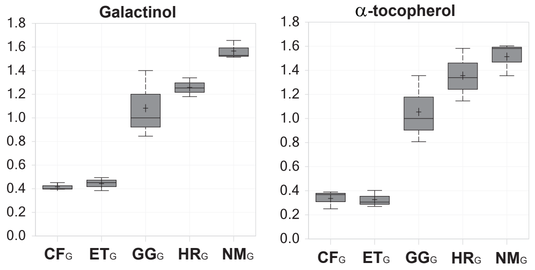


| SUPER_PATHWAY | CHEMICAL_NAME | PC1 |
|---|---|---|
| Amino Acid | tryptophan | -0.0745 |
| Amino Acid | kynurenate | -0.07372 |
| Amino Acid | phenylalanine | -0.07425 |
| Amino Acid | asparagine | -0.074 |
| Amino Acid | S-adenosylhomocysteine (SAH) | -0.07523 |
| Amino Acid | 3-methylhistidine | -0.07483 |
| Lipids | 1-palmitoyl-2-linoleoyl-GPC (16:0/18:2) | -0.0739 |
| Lipids | galactosylglycerol | -0.0741 |
| Cofactors | delta-tocopherol | -0.07421 |
| Secondary metabolism | phenylacetylglutamate | -0.07447 |

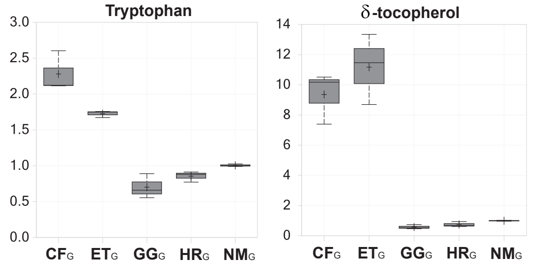


**PCA component 2 main drivers**

| SUPER_PATHWAY | CHEMICAL_NAME | PC2 |
|---|---|---|
| Amino Acid | 3-formylindole | 0.097938 |
| Amino Acid | methylsuccinate | 0.095242 |
| Lipids | 3-hydroxypalmitate | 0.084568 |
| Lipids | N,N,N-trimethyl-5-aminovalerate | 0.086186 |
| Lipids | 1-palmitoyl-2-(hydroxylinoleoyl)-GPC (16:0/18:2(OH))* | 0.08926 |
| Lipids | glycerophosphoglycerol | 0.08809 |
| Secondary metabolism | 2,4,6-trihydroxybenzoate | 0.091916 |
| Secondary metabolism | 4-hydroxycinnamate | 0.09933 |
| Secondary metabolism | ferulate | 0.087473 |
| Secondary metabolism | nicotianamine | 0.087503 |

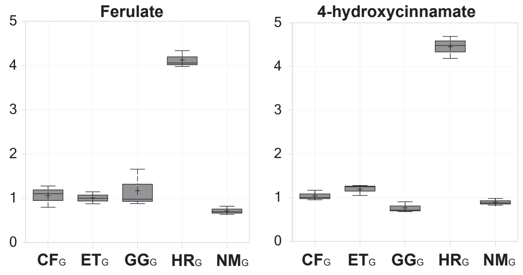


| SUPER_PATHWAY | CHEMICAL_NAME | PC2 |
|---|---|---|
| Amino Acid | fructosyllysine | -0.09474 |
| Amino Acid | stachydrine | -0.1014 |
| Carbohydrate | glucose | -0.09386 |
| Carbohydrate | arabitol/xylitol | -0.09663 |
| Lipids | 2-linoleoyl-GPE (18:2)* | -0.09127 |
| Nucleotide | N1-methyladenosine | -0.09504 |
| Nucleotide | cytosine | -0.09139 |
| Nucleotide | 2'-O-methyluridine | -0.09175 |
| Hormone metabolism | indole-3-carboxylate | -0.09004 |
| Secondary metabolism | 3-hydroxy-3-methylglutarate | -0.09029 |

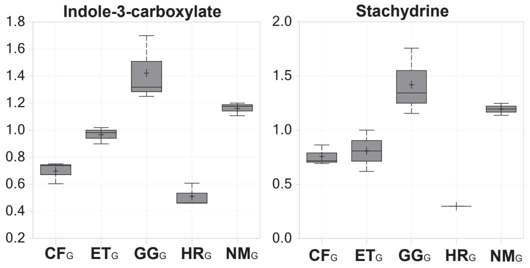


**Fig. S1. Compounds influencing separation along component 1 and component 2 of the grain principal component analysis (PCA) plot.** The compounds with the top 10 positive and top 10 negative scores are listed.
